# kSanity: A k-mer based application for precision bacterial strain detection and quantification

**DOI:** 10.1101/2025.09.04.674052

**Authors:** Michael France, Issac Chaudry, Joseph Elsherbini, Jacques Ravel

## Abstract

**Motivation:** Accurate detection and quantification of bacterial strains in clinical samples is necessary to measure their colonization and persistence. Past methods to achieve this relied either on strain-specific qPCR assays, or shotgun metagenomic read mapping approaches. The resident microbial community is a major source of interference in both assays because it can contain conspecific strains bearing similarity to the focal strain(s).

**Results:** We present kSanity, a k-mer based application for the detection and quantification of targeted bacterial strains in shotgun metagenomic data. Because kSanity uses exact string matches between the reads and reference, it is less sensitive to interference by conspecific strains. We test the performance of kSanity using a combination of *in silico* spike-in experiments, and *in vivo* observational data. Our results demonstrate that kSanity provides precise and accurate quantification of targeted bacterial strains, even when they are present at low sequence coverage in the metagenome.

**Availability and implementation:** kSanity is available at: https://github.com/ravel-lab/kSanity

## Introduction

Bacterial live-biotherapeutics are being increasingly explored as treatment options for a variety of health conditions (1). Formal clinical trials are needed to demonstrate their safety and efficacy and to establish their dosing regime. Failure of the live-biotherapeutic in such a trial could be driven by an inability of the bacterial strain(s) to engraft in the resident community (2). Characterizing the colonization and persistence of administered bacterial strains *in vivo* is therefore of critical importance in understanding their treatment efficacy. Tracking the abundance of a bacterial strain in mixed communities with a high degree of specificity is not a trivial task. Conspecific strains are naturally present in the microbiota of the treated population and can exhibit similarity to the focal strain thereby interfering with detection and quantification. In the past, qPCR assays, targeting regions thought to be specifical to a focal strain, have been used but the method is prone to false positives due to the presence of the targeted genomic region in conspecific strains or even other species (3, 4). Shotgun metagenomics offers a more attractive method as it enables the utilization of the focal strain’s whole genome and provides a characterization of the entire community. Yet disentangling the sequences originating from the focal strain from those of conspecific strains can prove troublesome due to high intrinsic genomic similarities and the shortcomings of read mapping software(5–7). Sequence read mapping software has been designed to maximize the number of reads mapping in order to accommodate for minor differences between the reads and the reference. While this approach is suitable for many applications (e.g. community composition profiling (8), RNAseq (9)), it complicates their use in precision strain tracking. Here, we present kSanity, a sequence k-mer based approach for detecting and quantifying bacterial strain(s) in shotgun metagenomic data. Sequence k-mers are short substrings of equal length that allow for exact matching between read and reference (10). We demonstrate that kSanity provides precision detection and quantification of bacterial strains in shotgun metagenomic datasets.

## Implementation

An illustration of the kSanity workflow can be found in Fig. 1 and includes two stages. In the first stage a reference database of unique and shared k-mers is constructed for the focal strain. In the second stage, a shotgun metagenomic dataset is split into k-mers and matched against the reference k-mers to detect and quantify the abundance of the focal strain.

**Fig. 1.**
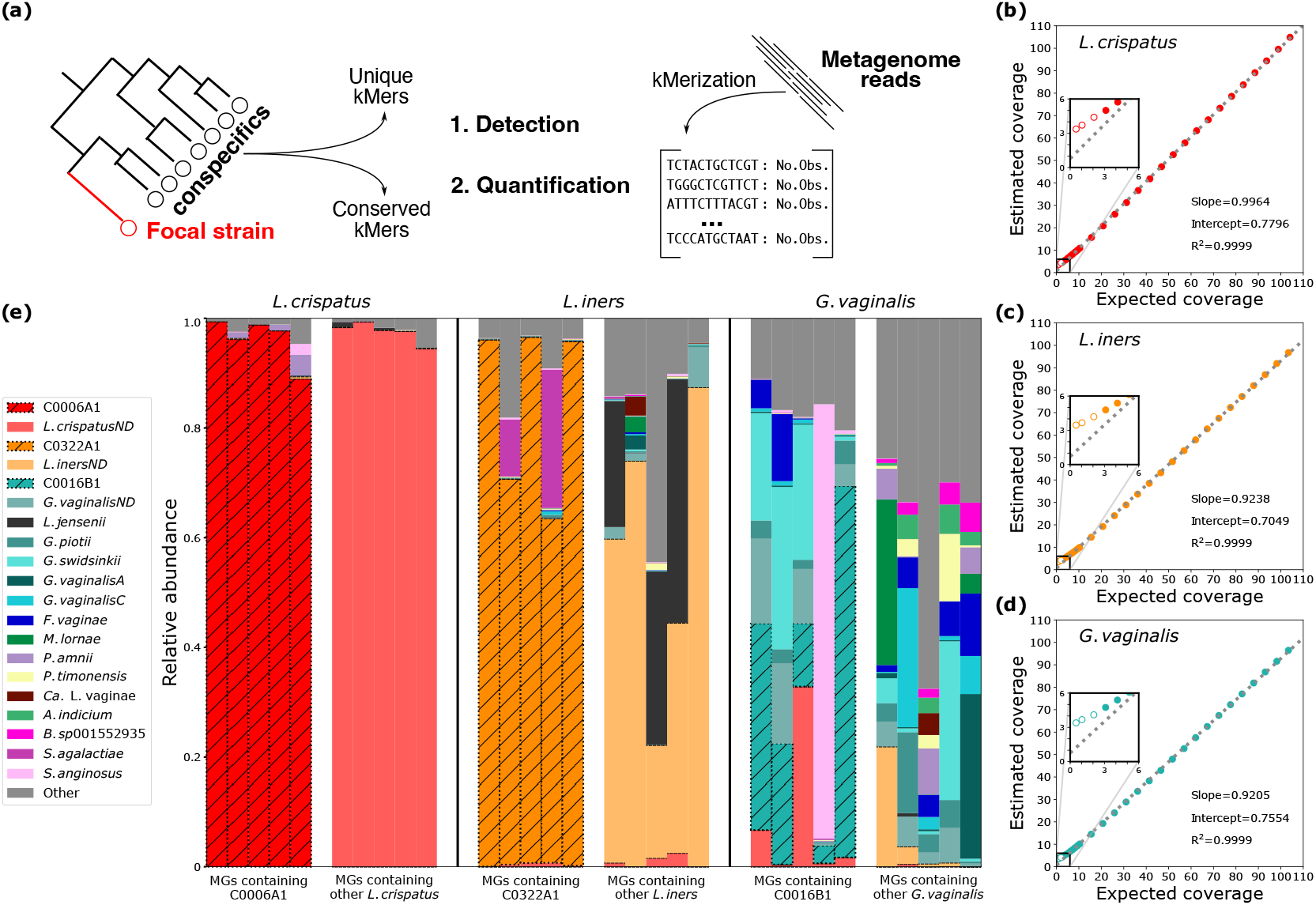
Schematic overview of kSanity (**a**). Results of *in silico* spike-in experiments for *L. crispatus* (**b**), *L. iners* (**c**), and *G. vaginalis* (**d**). Varying numbers of simulated sequence reads from target strains were added to real metagenomes that already contained the focal species. Coverage of the spiked strain was estimated using kSanity and regressed against the expected value. Open points represent samples where the percentage of strain-specific k-mers detected in the data was less than 50%. Inset plots zoom in on the region from 0-6X coverage to highlight the performance of kSanity at low read depths. Detection and quantification of target strains *in vivo* (**e**). For each strain (*L. crispatus*: C0006A1, *L. iners*: C0322A1, *G. vaginalis*: C0016B1), five metagenomes describing the vaginal microbiome of the participant from which the strain was isolated were selected, along with five metagenomes from other participants with a microbiome containing the same species. Stacked bar plots represent the taxonomic composition, with the relative abundance of the focal strain highlighted by thatched lines, as estimated by kSanity. The relative abundances of other *L. crispatus, L. iners*, and *G. vaginalis* strains are reported as not-differentiated (ND).

### Building the reference

In the first stage, a reference database of strain-specific and conserved k-mers is constructed using the whole genome sequences of the focal strain and conspecific strains (n>10). Each genome is split into k-mers of length *k*, with a 1bp sliding window, in both the forward and reverse directions. Exact string matches are then used to define which k-mers are unique to each strain’s genome (strain-specific) and which k-mers are shared among all of the strains (conserved k-mers). The k-mers specific to the focal strain are considered its “signature” and are used to estimate its coverage in the metagenome. The conserved k-mers are used to estimate the overall sequence coverage of the species in the metagenome which is used in the calculation of the focal strain’s relative abundance in the population.

### Detection and Quantification

In the second stage, sequence reads from a shotgun metagenome are split in k-mers of the same length used in the construction of the reference database, again with a 1bp sliding window. The number of times each k-mer is observed in the metagenome is recorded. Exact string matches are then made between the k-mers identified in the sequence reads and those in the reference database. A strain is considered detected if more than 50% of its strain-specific k-mers are identified in the metagenome, although this threshold can be adjusted depending on the use-case. Sequence coverage of the focal strain, and the species overall, is then determined using the average number of observances of the strain-specific and conserved k-mers, respectively. Two adjustments must be made to convert the k-mer estimates of coverage to be on the same scale as traditional read mapping based estimates. First, the k-mer value must be multiplied by two, because in k-mer space the length of the genome is doubled by the inclusion of both the forward and reverse directions. Second, the values must be adjusted for the lower contribution of a read to k-mer based coverage by multiplying the value by 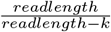. The abundance of the focal strain relative to that of the species is then calculated as the ratio of the average coverage of strain-specific and conserved k-mers.

## Examples

To validate kSanity and demonstrate its ability to detect and quantify bacterial strains, we performed tests using three different species prevalent in the human vaginal microbiome: *Lactobacillus crispatus, Lactobacillus iners*, and *Gardnerella vaginalis*. For each species we selected a focal strain with a complete genome sequence and then gathered publicly available whole genome sequences from conspecific strains (accession numbers in Supplementary Table 1). Each species genome dataset was reduced using dRep dereplicate function (11) (v3.1.1, -S algorithm=fastANI -SkipMash - sa=0.999 -clusterAlg=ward) to ensure that no two genome sequences were more than 99.9% identical in sequence. For each species, we performed tests using an *in silico* spike-in approach, and *in vivo* observational data. Tests were conducted using eight different k-mer lengths: 31, 37, 43, 49, 55, 61, 67, and 73. Performance was not sensitive to k-mer length (Supplementary Figure 1), and all reported results use a k-mer length of 55bp.

### *In silico* spike-in experiments

We first performed an *in silico* spike-in test to demonstrate kSanity could detect and precisely quantify bacterial strains. First, Illumina sequence reads of the focal strain were generated using inSilicoSeq (12) (v2.0.1, -coverage=uniform, -gc-bias, -model=novaseq). Varying numbers of the simulated reads (range: 10^4^ − 10^6^) were then added to a metagenome known to already contain that species (e.g. simulated *L. crispatus* were added to a *L. crispatus*-dominated metagenome). Detection of each strain (≥50% strain-specific k-mers) was achieved at around 3X coverage for all three species (≈50,000 reads). This enables kSanity to detect a bacterial strain even when it is at low relative abundance in the community, given sufficient sequencing depth. Next we estimated the sequence coverage of the spiked-in strain and compared it to the expected value based on the number of reads added (*L. crispatus*-Fig. 1b, *L. iners*-Fig. 1c, *G. vaginalis*-Fig. 1d). Linear regression lines were fit to the data using scipy (13) (v1.13.1, default settings), revealing the precision (r^2^>0.9999) and accuracy (slope>0.9 and intercept <0.8 for all species). This result demonstrates that kSanity provides rigorous detection and quantification of the relative abundance of a bacterial strain, even in the background of a metagenome composed of conspecific strains.

### *In vivo* observational data

In our second example, we sought to demonstrate that kSanity could detect and quantify a bacterial strain *in vivo*. To do this, we leveraged a previously published dataset of shotgun metagenomes describing the composition of the vaginal microbiome of participants (n=3) from which we had cultivated strains of *L. crispatus, L. iners*, or *G. vaginalis* (14, 15). Each participant had five longitudinal metagenomes available that were derived from samples collected ≈2 weeks apart. This dataset was further supplemented with vaginal metagenomes from other participants in the study that contained either *L. crispatus* (n=5), *L. iners* (n=5) or *G*.*vaginalis* (n=5). We used kSanity to detect and quantify the focal bacterial strains in all of the metagenomes, and then characterized the overall taxonomic composition of the communities using VIRGO2 (16) (v1, default settings). For each species, kSanity detected the focal strain only in the metagenomes derived from the corresponding participant from which the strain had been isolated, and not those derived from other participants (Fig. 1e). The estimated relative abundances of the focal *L. crispatus* and *L. iners* strains were above 90% and 60%, respectively at all five time points. However, the focal *G. vaginalis* strain fluctuated in relative abundance across the samples, ranging from comprising <5% to >60% of the community. This result demonstrates that kSanity maintains its performance when used to analyze metagenomes derived from clinical samples and is capable of quantifying the relative abundance of even uncommon strains in mixed communities.

## Conclusion

We presented kSanity, a software package for the detection and quantification of bacterial strains in shotgun metagenomic data. The k-mer based approach enforces exact string matches between sequence reads and reference genomes, thereby minimizing the risk for false-positive strain detections. In our example analyses, we found kSanity to provide a high degree of specificity in detection and accuracy in quantification. Although all testing was performed only with short-read metagenomes, reads from long read sequencing should also be suitable. We anticipate that kSanity will prove useful in the determination of colonization and persistence of live-biotherapeutic bacterial strains in clinical trials. However, kSanity also has applications in other settings where tracking a bacterial strain in a community is needed. For example, kSanity could be used to verify the strain composition of a bacterial product without requiring isolation. The software is implemented in Python 3, and is available on GitHub (https://github.com/ravel-lab/kSanity).

## Author Contributions

M.F., I.C., and J.R. conceived the approach, M.F. and J. E. conducted and analyzed the experiment(s), M.F., I.C., J.E., and J.R. wrote and reviewed the manuscript.

## Supplementary Data

**Supplementary Table 1**: Accession numbers of whole genome sequences and shotgun metagenomes used in the analysis

**Supplementary Figure 1**: Linear regression lines relating the expected and estimated coverages for different values of k. The bold line represents the value of k used in all of the presented analyses (k=55).

## Bibliography

1. T. M. G. Dronkers, A. C. Ouwehand, and G. T. Rijkers. Global analysis of clinical trials with probiotics. Heliyon, 6(7):e04467, 2020. doi: 10.1016/j.heliyon.2020.e04467.

2. Niv Zmora, Gili Zilberman-Schapira, Jotham Suez, Uria Mor, Mally Dori-Bachash, Stavros Bashiardes, Eran Kotler, Maya Zur, Dana Regev-Lehavi, Rotem Ben-Zeev Brik, Sara Federici, Yotam Cohen, Raquel Linevsky, Daphna Rothschild, Andreas E. Moor, Shani Ben-Moshe, Alon Harmelin, Shalev Itzkovitz, Nitsan Maharshak, Oren Shibolet, Hagit Shapiro, Meirav Pevsner-Fischer, Itai Sharon, Zamir Halpern, Eran Segal, and Eran Elinav. Personalized gut mucosal colonization resistance to empiric probiotics is associated with unique host and microbiome features. Cell, 174(6):1388–1405.e21, 2018. doi: 10.1016/j.cell.2018.08.041.

3. P. Kralik and M. Ricchi. A basic guide to real time pcr in microbial diagnostics: Definitions, parameters, and everything. Front Microbiol, 8:108, 2017. doi: 10.3389/fmicb.2017.00108.

4. C. R. Cohen, M. R. Wierzbicki, A. L. French, S. Morris, S. Newmann, H. Reno, L. Green, S. Miller, J. Powell, T. Parks, and A. Hemmerling. Randomized trial of lactin-v to prevent recurrence of bacterial vaginosis. N Engl J Med, 382(20):1906–1915, 2020. doi: 10.1056/NEJMoa1915254.

5. D. Albanese and C. Donati. Strain profiling and epidemiology of bacterial species from metagenomic sequencing. Nat Commun, 8(1):2260, 2017. doi: 10.1038/s41467-017-02209-5.

6. B. J. Smith, X. Li, Z. J. Shi, A. Abate, and K. S. Pollard. Scalable microbial strain inference in metagenomic data using strainfacts. Front Bioinform, 2:867386, 2022. doi: 10.3389/fbinf.2022.867386.

7. D. T. Truong, A. Tett, E. Pasolli, C. Huttenhower, and N. Segata. Microbial strain-level population structure and genetic diversity from metagenomes. Genome Res, 27(4):626– 638, 2017. doi: 10.1101/gr.216242.116.

8. A. Blanco-Miguez, F. Beghini, F. Cumbo, L. J. McIver, K. N. Thompson, M. Zolfo, P. Manghi, L. Dubois, K. D. Huang, A. M. Thomas, W. A. Nickols, G. Piccinno, E. Piperni, M. Puncochar, M. Valles-Colomer, A. Tett, F. Giordano, R. Davies, J. Wolf, S. E. Berry, T. D. Spector, E. A. Franzosa, E. Pasolli, F. Asnicar, C. Huttenhower, and N. Segata. Extending and improving metagenomic taxonomic profiling with uncharacterized species using metaphlan 4. Nat Biotechnol, 41(11):1633–1644, 2023. doi: 10.1038/s41587-023-01688-w.

9. D. Kim, J. M. Paggi, C. Park, C. Bennett, and S. L. Salzberg. Graph-based genome alignment and genotyping with hisat2 and hisat-genotype. Nat Biotechnol, 37(8):907–915, 2019. doi: 10.1038/s41587-019-0201-4.

10. C. Moeckel, M. Mareboina, M. A. Konnaris, C. S. Y. Chan, I. Mouratidis, A. Montgomery, N. Chantzi, G. A. Pavlopoulos, and I. Georgakopoulos-Soares. A survey of k-mer methods and applications in bioinformatics. Comput Struct Biotechnol J, 23:2289–2303, 2024. doi: 10.1016/j.csbj.2024.05.025.

11. M. R. Olm, C. T. Brown, B. Brooks, and J. F. Banfield. drep: a tool for fast and accurate genomic comparisons that enables improved genome recovery from metagenomes through de-replication. ISME J, 11(12):2864–2868, 2017. doi: 10.1038/ismej.2017.126.

12. H. Gourle, O. Karlsson-Lindsjo, J. Hayer, and E. Bongcam-Rudloff. Simulating illumina metagenomic data with insilicoseq. Bioinformatics, 35(3):521–522, 2019. doi: 10.1093/bioinformatics/bty630.

13. Pauli Virtanen, Ralf Gommers, Travis E. Oliphant, Matt Haberland, Tyler Reddy, David Cournapeau, Evgeni Burovski, Pearu Peterson, Warren Weckesser, Jonathan Bright, Stéfan J. van der Walt, Matthew Brett, Joshua Wilson, K. Jarrod Millman, Nikolay Mayorov, Andrew R. J. Nelson, Eric Jones, Robert Kern, Eric Larson,J J Carey, İlhan Polat, Yu Feng, Eric W. Moore, Jake VanderPlas, Denis Laxalde, Josef Perktold, Robert Cimrman, Ian Henriksen, E. A. Quintero, Charles R. Harris, Anne M. Archibald, Antônio H. Ribeiro, Fabian Pedregosa, Paul van Mulbregt, and SciPy 1.0 Contributors. SciPy 1.0: Fundamental Algorithms for Scientific Computing in Python. Nature Methods, 17:261–272, 2020. doi: 10.1038/s41592-019-0686-2.

14. M. France, L. Fu, L. Rutt, H. Yang, M. Humphrys, S. Narina, P. Gajer, B. Ma,J J Forney, and J. Ravel. Insight into the ecology of vaginal bacteria through integrative analyses of metagenomic and metatranscriptomic data. SRA, 2022. doi: PRJNA797778.

15. M. T. France, L. Fu, L. Rutt, H. Yang, M. S. Humphrys, S. Narina, P. M. Gajer, B. Ma, L. J. Forney, and J. Ravel. Insight into the ecology of vaginal bacteria through integrative analyses of metagenomic and metatranscriptomic data. Genome Biol, 23(1):66, 2022. doi: 10.1186/s13059-022-02635-9.

16. M. T. France, I. Chaudry, L. Rutt, M. Quain, B. Shirtliff, E. McComb, A. Maros, M. Alizadeh, F. A. Hussain, M. A. Elovitz, D. A. Relman, A. Rahman, R. M. Brotman, J. Price, M. Kassaro, J. B. Holm, B. Ma, and J. Ravel. Virgo2: Unveiling the functional and ecological complexity of the vaginal microbiome with an enhanced non-redundant gene catalog. bioRxiv, 2025. doi: 10.1101/2025.03.04.641479.

