## Supplementary figures and images for "kSanity: A k-mer based application for precision bacterial strain detection and quantification"

### Supplementary Figure 1

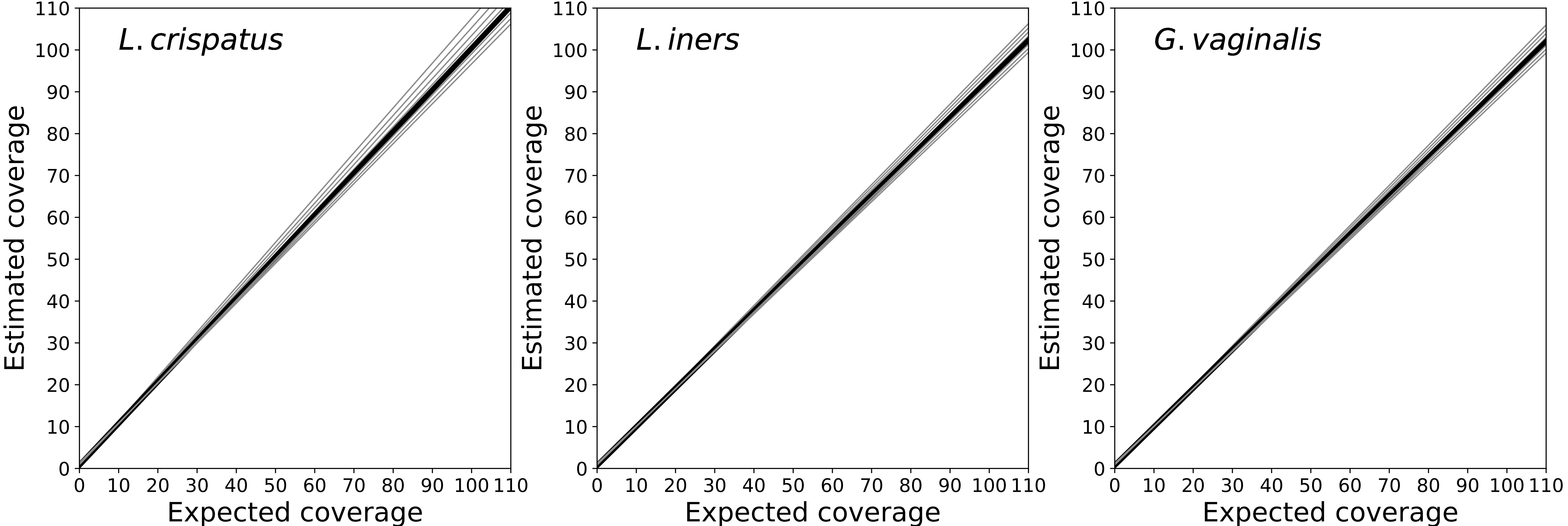
